# Testing the impact of gut microbiome rewilding on the health of captive bumble bees

**DOI:** 10.64898/2026.11.23.768741

**Authors:** Kristal M. Watrous, McKenna J. Larson, Tobin J. Hammer

## Abstract

Animals and plants are often raised in captivity for food, companionship, conservation, or research purposes, a practice which typically reduces the diversity of their microbiomes. This loss of microbial diversity may in turn compromise host nutrition, pathogen resistance, and reproduction. Microbiome “rewilding”—restoring microbes present in the wild, but lost in captivity—has been proposed as a way to ameliorate these effects. However, it has only been tested in a handful of taxa. We evaluated the efficacy of microbiome rewilding in bumble bees (*Bombus*), a group of ecologically and agriculturally important pollinators. In captivity, bumble bees retain host-specialized “core” gut bacteria, but lack the diverse environmental bacteria commonly present in the gut of wild bees. We inoculated captive *B. impatiens* colonies with five species of environmental bacteria isolated from wild *B. impatiens*. Rewilding did not affect bee survival, reproduction, or nutrition (glycogen or lipid levels). We conducted a second experiment with *Arsenophonus*, the environmental bacterium exhibiting the highest colonization rate in captive bees, to test its role in resistance against an opportunistic bacterial pathogen. While inoculation with gut material from captive bees (which is dominated by core bacteria) completely protected hosts from pathogen infection, *Arsenophonus* inoculation had no effect. These findings demonstrate that microbiome rewilding is not always effective for sustaining the health of captive hosts.

## Introduction

Animals and plants are raised in captivity for a variety of reasons, including food production, companionship, scientific research, and conservation (e.g., captive breeding programs). A common consequence of this practice is the loss of diverse microbial symbionts that are normally present in wild populations [1–4]. In turn, the loss of diversity could compromise microbiome functioning [5–7], perhaps contributing to the impaired health and reproduction seen in certain hosts when raised in captivity (e.g., [8,9]).

The restoration of microbes present in the wild, but lost in captivity, has been proposed as a way to support the health of captive hosts [3,10–14]. This “microbiome rewilding” approach has been successful in various animal and plant taxa [15–17]. However, there are also theoretical arguments questioning the effectiveness of microbiome rewilding. First, captivity entails unique diets, pathogens, and stressors not present in the wild, a context in which microbiome flexibility—including losses of microbes—may actually be advantageous [18–20]. Second, some wild microbes may be opportunistic pathogens, parasites, or commensals [21,22]. Third, rewilding generally requires microbes to be cultivated *in vitro* and delivered to hosts as probiotics. But cultured microbes may fail to colonize hosts, even if isolated from the same host species [23–25]. Given these challenges, the efficacy of microbiome rewilding should be evaluated on a host-by-host basis.

Bumble bees (Apidae: *Bombus*) are an important group of insect pollinators whose management and conservation might benefit from microbiome rewilding. Some species of bumble bees are in decline, likely due in part to disease [26–28]. For highly endangered bumble bee species, such as *B. affinis*, captive rearing may become an important tool for population recovery [29]. Certain other *Bombus* species, including *B. impatiens* in North America, have been domesticated and produced by commercial suppliers for crop pollination [30]. Success of microbiome rewilding in species like *B. impatiens* could prove useful for agriculture and could indicate promise for captive rearing of *Bombus* generally.

Bumble bee workers—which make up most of the individuals in a colony and whose gut microbiomes are best-studied—typically harbor host-specialized gut bacteria including *Snodgrassella*, *Gilliamella*, *Schmidhempelia*, *Bifidobacterium, Bombiscardovia*, *Lactobacillus*, and *Bombilactobacillus* [31,32]. This “core” microbiome has been shown to confer protection against pathogens including the trypanosomatid *Crithidia* spp. [33–35] and the bacterium *Serratia marcescens* [36]. However, wild worker bees may harbor a diverse array of environmental bacteria, sometimes at even higher abundances than the core bacteria (e.g., [37–39]. How these environmental bacteria affect bumble bee fitness is generally unknown, but effects may range from beneficial, to neutral, to pathogenic, depending on the species and context [36,40].

In captive bumble bee colonies reared indoors, worker gut microbiomes usually lack diverse environmental bacteria and are instead dominated by core bacteria [41–46]. Such colonies are generally maintained under climate-controlled conditions and fed with sugar syrup and honey bee-collected pollen, which may or may not be sterilized [30]. In contrast, workers from colonies that are bred or initiated indoors, but allowed to forage for pollen and nectar from flowers in the environment, can exhibit high levels of environmental bacteria [42,47–49]. This discrepancy suggests that the loss of environmental bacteria in indoor-restricted colonies is mainly driven by a lack of exposure to floral microbes, not to other aspects of the captivity environment or to the domestication process.

In theory, an absence of environmental bacteria could contribute to difficulties with captive rearing of bumble bees. Success rates for establishing colonies indoors from wild-caught queens vary greatly across species but are often quite low (0–54%), and some species have never been successfully reared [50]. The causes are not fully known, but one factor (among many) may be that queens lack cues in their diet or environment that induce them to produce brood [51]. In other hosts such as mosquitoes and tubeworms, environmental bacteria can stimulate development [52,53]. Further, pathogens such as *Crithidia* or *Nosema* are often present in commercially reared bumble bees [54–57], which could limit effectiveness for crop pollination and spread pathogens to wild bees. Low microbiome diversity has been shown to facilitate pathogen colonization in other hosts [6]. Therefore, restoring diverse environmental bacteria has the potential to help ameliorate these challenges.

In this study we used *Bombus impatiens* as a model to test whether the health of captive bumble bees can be improved by rewilding with environmental bacteria. We selected five environmental bacterial species belonging to different families, in order to maximize potential functional diversity. These bacteria were isolated from the gut of wild *Bombus impatiens* workers, wherein they can reach high relative abundances [36,58]. Cultured environmental bacteria were inoculated into commercially produced *B. impatiens* colonies via their food, similarly to how probiotics are delivered to honey bee colonies [59,60]. After an acquisition period in the colony, young workers were moved into cages (“microcolonies”) to measure survival, brood production, microbial colonization, and stores of two important nutrients. In a separate experiment testing impacts on pathogen resistance, gnotobiotic workers were exposed to a focal environmental bacterium and challenged with an opportunistic bacterial pathogen [36]. Overall, the findings illustrate challenges inherent to microbiome rewilding and raise questions about how much, and why, its efficacy varies across host taxa.

## Methods

### Bee rearing

We obtained eight commercial *Bombus impatiens* colonies (Beneficial Insectary, Biobest Group, CA, USA), and provided them with a feeder filled with a commonly used artificial nectar solution (50% sucrose with additives; [61]) and non-sterile, multifloral honey bee-collected pollen (Koppert, Howell, MI) ground and mixed with sugar solution as a pollen dough. Colonies were randomly assigned to either the rewilding (referred to as “inoculated” below) or control treatment. Beginning one day after receipt, each colony was inoculated once per week for nine weeks with either a mixture of the environmental bacteria (described below) or a control inoculum (1 mL PBS), pipetted onto a ball of pollen.

To track known-age worker bees, we removed cocoons containing worker pupae from each colony, placed them in a cage in an incubator (Invictus Drosophila incubator) at 35 °C, and checked them daily for emergence. Within 24 hours of emergence each worker was marked with a unique numbered tag and then returned to the natal colony.

### Inoculum preparation

Environmental bacteria were isolated from wild *Bombus impatiens* workers and identified by Sanger sequencing as described in Nelson et al. [36] and Watrous et al. [58]. To prepare the inoculum, five strains were selected representing five different bacterial families and cultured using the conditions listed in Table 1. For each strain individually, bacterial cells were harvested and suspended in phosphate-buffered saline (PBS) at an OD of 2.0. We then combined 200 µL of each strain, totaling 1 mL of solution, to create the inoculum for one colony.

**Table 1.** The five isolates of environmental bacteria, sourced from wild *Bombus impatiens*, that were used to test microbiome rewilding. Bacteria were cultured under ambient oxygen conditions unless otherwise noted.

| Bacterium (Isolate ID) | Family | Medium | Growth conditions | Time |
|---|---|---|---|---|
| *Acinetobacter* (BSJ) | Moraxellaceae | Nutrient agar | 30 ºC | 48 h |
| *Arsenophonus* (PJL) | Morganellaceae | Columbia blood agar | 30 ºC | 168 h |
| *Fructobacillus* (4C7) | Lactobacillaceae | MRS 2% fructose agar | 30 ºC | 48 h |
| *Pseudomonas* (6AN) | Pseudomonadaceae | BHI agar | 30 ºC | 24 h |
| *Rosenbergiella* | Enterobacteriaceae | Columbia blood agar | 35 ºC at 5% | 24 h |
| (T41) |  |  | CO<sub>2</sub> |  |

### Fitness and nutrition experiment

Tagged workers were left in their natal colony for 5-6 days, the typical timeframe needed for newly emerged *B. impatiens* workers to acquire their core microbiome [43]. After this acquisition period, workers were removed from the natal colony and placed in microcolonies (i.e., small cages without the queen) of three workers each [62]. Each microcolony was maintained in a growth chamber at 27 °C and provided filter-sterilized 50% sucrose syrup and gamma-irradiated sterile multifloral honey bee-collected pollen patties *ad libitum*. In total, 3-6 microcolonies were established per natal colony.

To measure gut colonization and nutrient levels in the abdomen, half of the microcolonies from each natal colony were maintained for 14 days, after which we anesthetized the bees on ice and removed the midgut and hindgut. Each combined gut sample was h omogenized with a sterile pestle in 1 mL PBS then stored at −20 °C for later DNA extraction and 16S rRNA gene amplicon sequencing to characterize the bacterial community. Each bee body was stored at −20 °C for lipid and glycogen assays. To measure survival and reproduction, the other half of the microcolonies from each natal colony were fed and monitored for up to 56 days, with survival checked every 2-3 days. Reproductive output was quantified by counting the total number of eggs, larvae, pupae and adult males produced by each microcolony [36].

### Pathogen resistance experiment

We tested the ability of one of the five environmental bacteria, *Arsenophonus*, to protect bees against colonization by *Serratia marcescens*, a common opportunistic pathogen of *B. impatiens* [36]. We used three new commercial *B. impatiens* colonies from the same supplier as the previous experiment. Following an established approach for generating gnotobiotic bumble bee workers [36], we removed cocoons containing worker pupae, surface-sterilized them, and placed them in an incubator at 35° C. Upon adult emergence we placed three workers in each microcolony and assigned them to one of three treatments: core-inoculated (20 µL of fresh gut material from a pool of 4 workers from the matching natal colony, pestle-homogenized and suspended in 1 mL PBS; [34,35]; *Arsenophonus*-inoculated (20 µL *Arsenophonus* strain PJL at OD 2.0 in PBS); or axenic control (20 µL PBS only). Inoculum was applied to 2 g sterile pollen patties on the day of adult emergence, and bees remained in microcolonies kept at 27° C for four days to allow the inoculated bacteria to colonize the gut. On day five we inoculated all microcolonies with approximately 10^8^ colony-forming units (CFUs) of an overnight culture of a *Serratia marcescens* strain isolated from honey bees and labeled with a kanamycin resistance marker (originally described in [63]) in 20 μL PBS onto 2 g pollen patties. After another five days, guts were dissected and homogenized as described above. Gut homogenates were serially diluted and plated on LB agar with 20 µg/mL kanamycin with four technical replicates per sample. After 24 h incubation at 30 °C we counted CFUs of *Serratia* to estimate abundance per gut (2.5 × 10^2^ to 6 × 10^7^ CFUs).

### Nutrition assays

Bee abdomens were prepared for lipid and glycogen quantification following a modification of the protocol in [64]. Each abdomen from the fitness and nutrition experiment, frozen at −20 °C with gut already removed, was placed in an open 1.7 mL vial in a drying oven at 50 °C for two days to dry. Dry weight was recorded, then the abdomen was homogenized in a 2 mL vial with 900 µL molecular-grade water and one 5 mm chrome bead. Samples were homogenized using a Beadblaster 96 (Benchmark), shaking for 2 minutes at 1000 rpm, 1 minute rest, and 2 minutes at 1000 rpm, followed by centrifugation at 12700 rpm for 5 minutes.

From the homogenized sample, the liquid fraction was used for glycogen assays while the solid organic fraction was used to quantify lipids. To measure glycogen for each sample, 40 µL of liquid homogenate was mixed with 10 µL of 18% sodium sulfate and 700 µL of 1:1 chloroform:methanol solution. This mixture was centrifuged at 12700 rpm for 5 minutes. The supernatant was removed, leaving the glycogen as a precipitate. Residual chloroform was evaporated off, then the glycogen was resuspended in 500 µL of molecular water and vortexed. We added 150 µL of anthrone reagent (1 mg/mL of anthrone in concentrated sulfuric acid) to 50 µL of isolated glycogen sample. Assay plates were incubated at 4° C for 10 minutes, then 100° C at 300 rpm in a thermomixer (Eppendorf ThermoMixer C) for 20 minutes, then at room temperature in the dark for 20 minutes. Absorption values were measured at 620 nm (Molecular Devices SpectraMax M2e), blank subtracted, and compared to glucose standards (D-(+)-Glucose analytical standard, Sigma-Aldrich; 210, 180, 150, 120, 90, 60, 30 µg/mL).

Lipids were quantified using a sulfo-phosphovanillin assay used previously in [64]. To the homogenized organic solids we added 1000 µL of 2:1 chloroform:methanol solution then thoroughly vortexed the samples to dissolve lipids, followed by centrifugation for 2 minutes at 12700 rpm. The liquid portion of the sample was decanted into a 4 mL glass vial, pipetting around the solids. An additional 1000 µL 2:1 chloroform:methanol was added to each vial, then capped and placed in the refrigerator overnight to allow physical separation of layers. After overnight separation samples were stored at −80 °C. For each sample 50-100 µL was added in triplicate to wells of a 24-well microplate, then chloroform was evaporated off at 90 °C. Then 100 µL of concentrated sulfuric acid was added to each well and incubated at 90 °C for 20 minutes. Plates were cooled quickly to room temperature on an ice block, then background absorbance was measured at 540 nm (Molecular Devices SpectraMax M2e). We then added 50 µL of phosphovanillin solution (0.2 mg vanillin per 1 mL of 17% phosphoric acid) per well and allowed color to develop for 10 minutes in darkness before measuring final absorbance at 540 nm. After blank subtraction we subtracted sulfuric acid background absorbance from the phosphovanillin final absorbance and compared to cholesterol standards (Cholesterol solution, 10mg/mL in chloroform, Sigma-Aldrich; 80, 70, 60, 50, 40, 30 µg/mL)

### DNA sequencing

We extracted genomic DNA from frozen gut homogenates. We placed 250 µL of gut homogenate in a tube with ∼0.5 mL of 0.1 mm zirconia silica beads (BioSpec) and 750 µL ZymoBIOMICS Lysis Solution then homogenized with a tissue lyser (Benchmark Beadblaster96) at 1800 rpm for 2 minutes, left to rest for 1 minute at room temperature, then at 1800 rpm again for 2 minutes [36]. These lysed samples were stored at −70 °C until DNA extraction using the ZymoBIOMICS 96 DNA Kit. Following best practices [65], three extraction blanks and three technical replicates of ZymoBIOMICS microbial community standard were included.

Genomic DNA was PCR-amplified using barcoded universal primers (515F/806R) targeting the V4 region of the 16S rRNA gene [66]. PCR amplification methods followed those of [36]. Cleanup and normalization of PCR products was done using SequalPrep normalization plates. Libraries were pooled and sequenced on an Illumina MiSeq v2 (2 x 250), by the University of Colorado Boulder / Cooperative Institute for Research in Environmental Sciences’ Center for Microbial Exploration. Raw reads are available from NCBI (BioProject no. PRJNA1561584).

### Sequence data analysis

Amplicon sequences were processed and analyzed using the same protocol described in [67]. Briefly, cutadapt was used to remove adapters and primers from the demultiplexed data [68]. DADA2 was used to trim, quality-filter, and denoise data to yield amplicon sequence variants (ASVs) [69]. The RDP Naïve Bayesian Classifier [70] and SILVA 138.2 SSU rRNA reference database [71] were used to assign taxonomy to the ASVs. The ASV table was imported into R for further analysis with the package mctoolsr (https://github.com/leffj/mctoolsr). We removed very low-abundance ASVs (those with < 10 total reads across all samples), and those without domain-level classification. ASVs classified as mitochondria, chloroplast, or eukaryotic comprised an average of 0.57% (range 0 - 6.21%) of sequences across bee gut samples and were removed. We confirmed that the eight bacterial genera present in the ZymoBIOMICS microbial community standard comprised the top eight genera (in terms of mean relative abundance) in the respective sequence libraries. Putative reagent contaminant ASVs were identified and removed using decontam [72]. Bee gut samples that originally had >2% contaminant ASV sequences were discarded (removing two). Bee gut sequence libraries were randomly subsampled (rarefied) to an even sequence depth of 10,000 sequences, which removed one sample.

### Statistical Analyses

To examine the effect of exposure to environmental bacteria on bee survival, we constructed a Cox mixed-effects model with bee survival as the response with microbiome treatment as a fixed effect and natal colony as a random effect. To analyze bee glycogen levels, lipid levels, and total microcolony reproduction, we constructed linear mixed-effects models (‘LMER’) with microbiome treatment as a fixed effect and natal colony as a random effect. The response variables were µg glycogen per µg dry bee abdomen (*ln*-transformed), µg total lipids per 100 µL of sample per µg dry bee abdomen (ln-transformed), and total brood in each microcolony (*ln* + 1 transformed), respectively. To assess the effect of microbiome treatment on *Serratia* colonization, we constructed an LMER with *S. marcescens* CFU counts (*ln* + 1 transformed) as the response variable, microbiome treatment as a fixed effect and natal colony as a random effect.

Statistical analyses were conducted in R v4.5.0 [73] and RStudio v2026.4.0.526 [74]. For the Cox mixed-effects model we used the ‘coxme’ package [75]. LMERs were constructed using the ‘lmer()’ function in the package ‘lme4’ [76]. To assess the significance of fixed effects, we used Wald chi-square tests with type III sums of squares for LMERs using the Anova() function in the ‘car’ package [77].

## Results

Inoculating captive bumble bee (*Bombus impatiens*) colonies with environmental bacteria derived from wild conspecifics did not affect survival of focal worker bees, after accounting for natal colony as a random effect (Fig 1A; Cox mixed-effects model, p = 0.75). Environmental bacteria also had no significant effect on worker reproduction (Fig 1B; LMER: χ^2^ = 1.057, df = 1, p = 0.3039), again after accounting for natal colony as a random effect.

**Figure 1.**
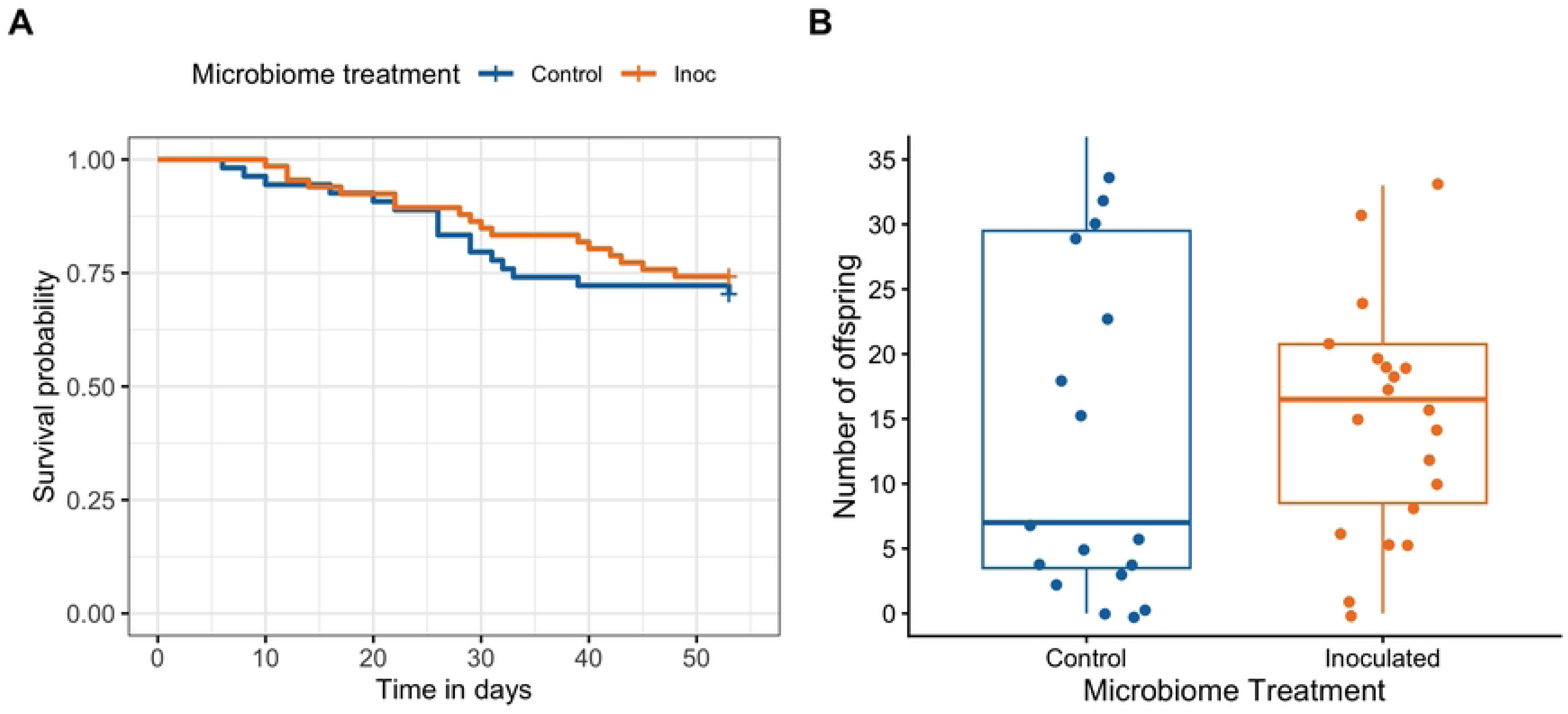
Survival and reproduction of captive-reared worker bees exposed to either environmental bacteria (“inoculated”) or sterile buffer only (“control”). Bees were sourced from four natal colonies per treatment. (A) Inoculation with environmental bacteria does not alter survival. (B) Inoculation does not alter the total production of offspring, including eggs, larvae, pupae, and adults present at microcolony termination (56 days).

A separate cohort of 14-day-old worker bees, from the same natal colonies, was used to assess gut colonization by environmental bacteria. Overall, gut microbiomes were dominated by core (i.e. bumble bee-specialized) bacterial genera (Fig 2). Of the five strains of environmental bacteria that were cultured and inoculated into colonies (Table 1), only *Arsenophonus* (corresponding to strain PJL) exhibited a mean relative abundance (across all individual bees) above 0.1% of 16S rRNA gene sequences. *Arsenophonus* ranged from 0-98% of sequences, with mean (± SEM) values of 0.39% ± 0.21% in control bees and 5.29% ± 2.49% in inoculated bees (Fig 2). The other four genera listed in Table 1 were either completely absent or very rare, with combined relative abundances of 0.013% ± 0.0031% (mean ± SEM) in control bees and 0.016% ± 0.0052% in inoculated bees.

**Figure 2.**
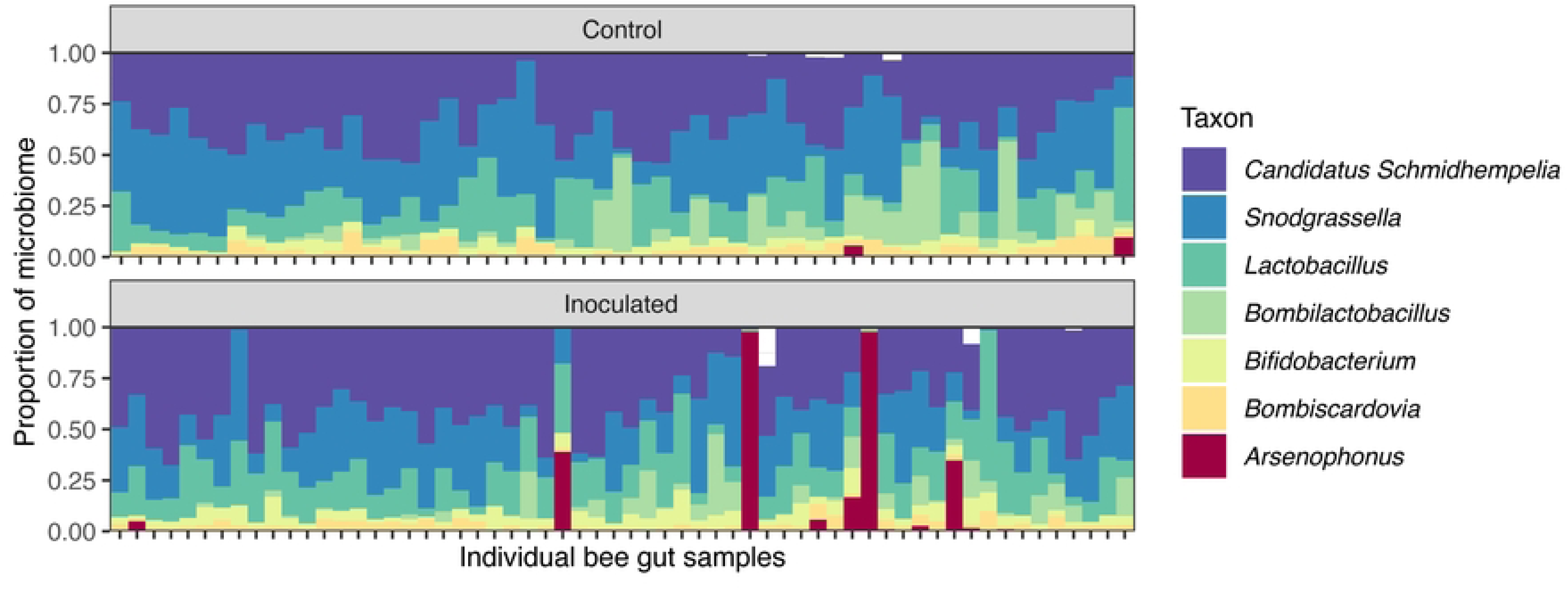
Gut microbiome composition in 14-day-old worker bees exposed to either environmental bacteria (“inoculated”) or sterile buffer only (“control”). Each vertical bar is one bee, and the y-axis shows proportion of 16S rRNA gene sequences classified as a given bacterial genus. Of the five environmental bacterial taxa in the inoculum, only *Arsenophonus* was detected in bee guts. The other six genera in the legend are core (i.e., bumble bee-specialized) bacteria. All genera with > 0.1% mean relative abundance across the sample set are shown, with other taxa represented by white space.

The same 14-day-old workers as described above were also evaluated for abdominal lipid and glycogen levels. Exposure to environmental bacteria had no significant effect on total lipids (Fig 3A; LMER: χ^2^ = 0.7552, df = 1, p = 0.3848), or glycogen (Fig 3B; LMER: χ^2^ = 3.457, df = 1, p = 0.0629) after accounting for natal colony as a random effect.

**Figure 3.**
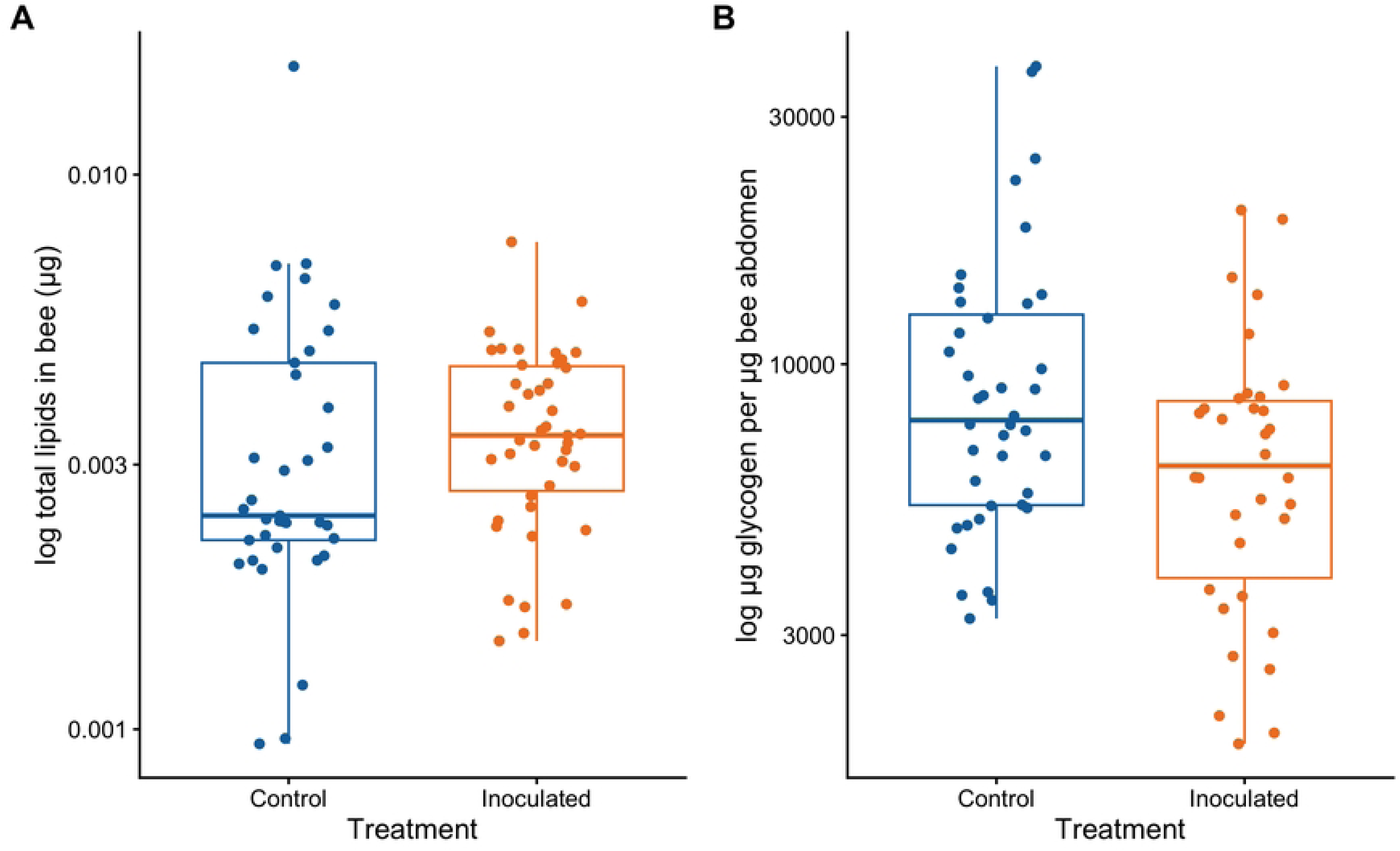
Levels of abdominal lipids and glycogen in 14-day-old worker bees exposed to either environmental bacteria (“inoculated”) or sterile buffer only (“control”). Neither lipids (A) nor glycogen (B) significantly differed between treatments.

A separate experiment, with gnotobiotic worker bees from three new natal colonies of *B. impatiens*, was conducted to test the effect of captive versus environmental (i.e. *Arsenophonus*) bacteria on resistance to the opportunistic pathogen *Serratia marcescens*. Colonization by *S. marcescens* differed strongly among the three treatment groups (Fig 4; LMER: χ^2^ = 156.87, df = 2, p << 0.001). Exposure to captive-derived bacteria almost completely prevented *S. marcescens* colonization (median 0 CFUs) relative to axenic bees, which harbored a median of 1 * 10^5^ CFUs (Fig 4; Wilcoxon rank sum test with Bonferroni adjustment: p << 0.001). However, *Serratia* colonization was not affected by exposure to *Arsenophonus*, with a median of 1.39 * 10^5^ CFUs (Fig 4; Wilcoxon rank sum test with Bonferroni adjustment: p = 1).

**Figure 4.**
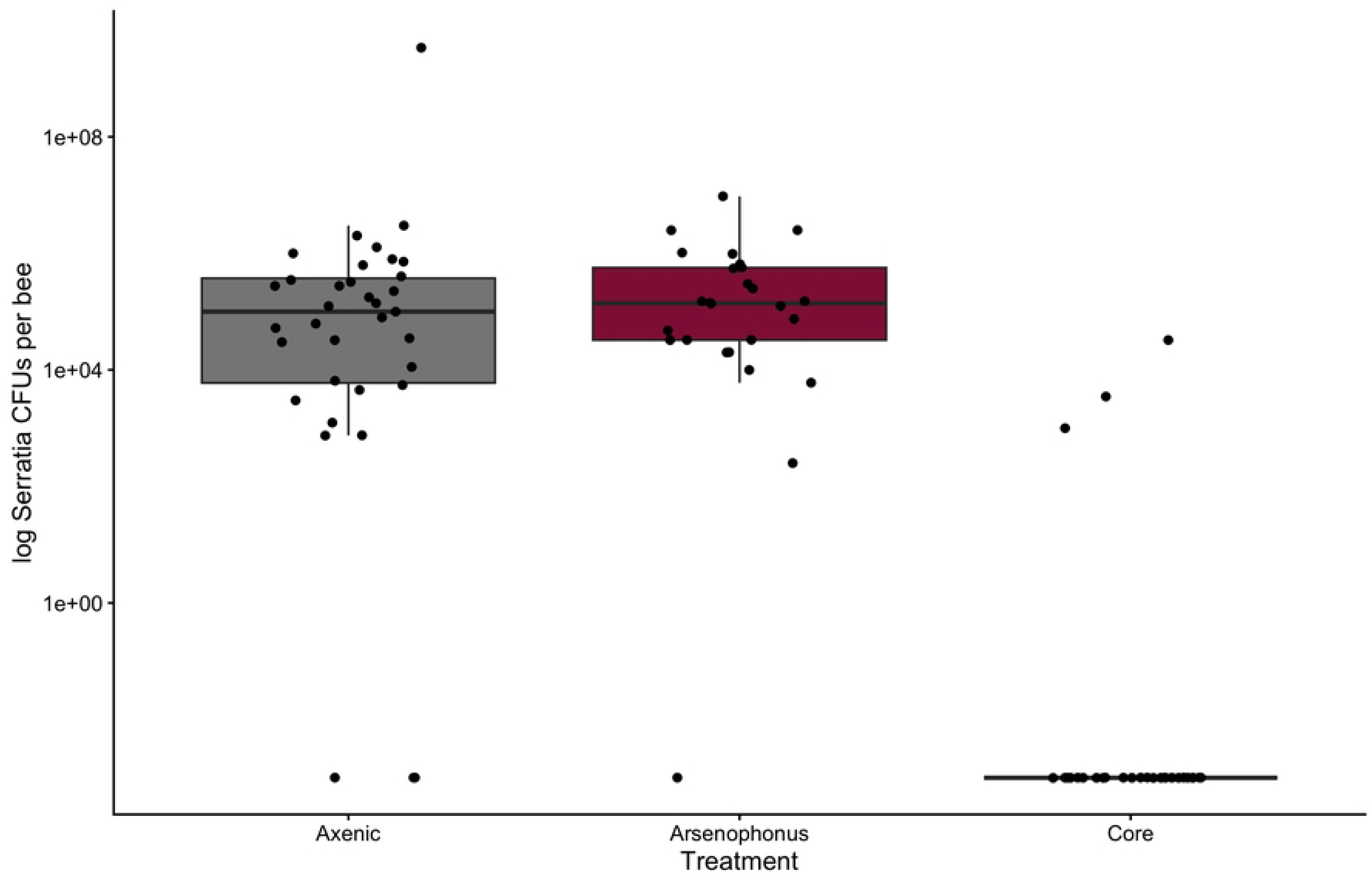
Microbiome effects on gut colonization by an opportunistic bacterial pathogen (*Serratia marcescens*) in gnotobiotic worker bees. Axenic workers were initially exposed to either the environmental bacterium *Arsenophonus*, gut material from unmanipulated nestmates (presumed to be dominated by core bacteria; “Core”), or sterile buffer only (“Axenic”). *Serratia* colonization is strongly reduced in the Core, but not *Arsenophonus* treatment.

## Discussion

We tested the impact of microbiome rewilding on the health of captive bumble bees. Five strains of environmental bacteria, isolated from the gut of wild *Bombus impatiens*, were cultured and inoculated into captive *Bombus impatiens* colonies. Overall, despite the much-touted promise of microbiome rewilding (e.g., [10,11,16]), we found no impact on any of the health and fitness metrics that we analyzed. Focal age-controlled bees inoculated with environmental bacteria in their colonies exhibited similar survival rates, brood production, and levels of two important classes of nutrients (lipids and glycogen), as compared with control bees. The values of these metrics were similar to previous studies of captive *B. impatiens*, suggesting that our protocols are reliable [36,78].

The only effect of rewilding close to statistical significance was with glycogen. However, our data suggest that any such effect is likely to be negative, not positive. Some of the environmental bacteria may induce a host immune response, increasing metabolic rates and consumption of glucose, the building block of glycogen [79,80]. *Arsenophonus* is a likely candidate, given that it was the only environmental bacterium to successfully (albeit sporadically) colonize *B. impatiens*. Further, bumble bee-associated *Arsenophonus* encodes numerous virulence factors and toxins in its genome [40], and in honey bees, *Arsenophonus* is associated with host mortality [81].

The general lack of gut colonization by the environmental bacteria may, at least in part, explain the lack of impact on host phenotypes. Poor colonization was surprising because the chosen bacterial strains were isolated from the gut of wild conspecific hosts in which the corresponding genera can reach high relative abundances [36]. Further, the bacteria were applied to the sole pollen source for each colony in high doses, ensuring that bees would be exposed. Lack of colonization by environmental bacteria could be due to resistance from core bacteria in the colony, as indicated by our *Serratia marcescens* data (Fig. 4) and previous work [36]. Additionally, changes in physiology during *in vitro* cultivation [82] may have rendered the environmental bacteria less capable of host colonization. Both mechanisms could pose barriers to microbiome rewilding or similar practices (such as probiotic administration) in bumble bees.

In a second experiment designed to test for pathogen protection, we did not find any effect of inoculation with a focal environmental bacterium (*Arsenophonus* strain PJL) on infection by the bacterial pathogen *Serratia marcescens*. This lack of effect could, in theory, again be explained by a lack of *Arsenophonus* colonization. However, *Arsenophonus* successfully colonized some bees in the first experiment, and bees in the second experiment were axenic prior to inoculation, which could be expected to facilitate colonization. In contrast, we observed that inoculation with gut material from captive nestmates completely protected bees from *Serratia* infection, in agreement with previous results [36]. Such material was almost certainly dominated by core bacteria, based on our 16S rRNA gene sequencing data (Fig. 2) and previous work with captive *B. impatiens* [43,45].

Certain caveats should be considered. First, variation in the measured phenotypes between natal colonies may have obscured subtler treatment effects, especially with low colony-level replication. In bumble bees, different colonies—even from the same supplier or wild population—often maintain distinct microbiomes in the laboratory [43,47,83]. This microbiome variation, in addition to host genetic variation, may drive differences in survival, reproduction, and other parameters, reinforcing the importance of including replicate colonies in bee experiments and statistical analyses. Second, experiments were conducted under stable and optimal conditions. Thus, potential effects of rewilding on host stress resistance or resilience were not tested. Third, there are many more species of environmental bacteria associated with wild *B. impatiens* than the five tested here [36,84]. Genomic analysis may be useful in prioritizing strains for testing, based on traits such as metabolic potential or host colonization factors.

Wild strains of core bacteria may be a promising alternative to environmental bacteria for rewilding the bumble bee gut microbiome, but come with their own challenges. Because they are already adapted for growth in the bee gut, core bacteria may colonize captive bees more readily [60]. Further, they appear to have consistent benefits for resistance to pathogen infection, as discussed above. On the other hand, microbe-by-host-genotype interactions [24] and competition with captive-associated core bacteria occupying similar niches—e.g., through priority effects [85] and antagonism [86]—may constrain the establishment of wild core strains. Future studies should investigate the potential benefits, as well as limitations, of using core bacteria for bumble bee rewilding or probiotic development.

Given the enormous variation in microbiome form and function across host taxa [87], it can be expected that the efficacy of rewilding will also vary. For example, rewilding may be particularly impactful for plants and animals which horizontally acquire their microbiome from the environment [88]. Such hosts would lose most of their microbiome diversity when brought into a captive rearing environment, creating a need for rewilding. In contrast, hosts that vertically transmit microbes, such as social bees, should maintain a greater portion of their wild microbiome in captivity without any intervention necessary (assuming parent to offspring transmission is not disrupted). In these hosts, the microbes that are present in the wild, but lost in captivity, could be largely commensal or pathogenic. Perhaps the “shedding” of such microbes is beneficial, and their reintroduction, through rewilding, could actually be harmful.

Although our findings indicate that the latter scenario may apply to bumble bees, more bacterial strains and species will need to be tested, under more conditions, before definitive conclusions can be drawn. We suggest that a promising avenue for future work is to model combinations of host and microbial traits to predict the likelihood of microbiome rewilding success, which can then be tested with experiments.

## Acknowledgments

We thank the University of Colorado Boulder / Cooperative Institute for Research in Environmental Sciences’ Center for Microbial Exploration for their sequencing services, and members of the Hammer Lab at UC Irvine for feedback. We have not used AI-assisted technologies in creating this article.

## Data availability

Raw sequence data have been uploaded to the European Nucleotide Archive (PRJNA1561584). All other data and R scripts are available from the Dryad Digital Repository: https://doi.org/10.5061/dryad.zgmsbccvz

## Ethics

This work did not require ethical approval from an animal welfare committee.

## Author contributions

Conceptualization: TJH, KMW. Data Curation: KMW, MJL, TJH. Formal Analysis: KMW, TJH. Funding Acquisition: TJH. Investigation: KMW, MJL. Visualization: KMW, TJH. Writing – Original draft preparation: KMW, TJH. Writing – Review & Editing: KMW, MJL, TJH.

